# Spectral estimation of *in-vivo* wheat chlorophyll a/b ratio under contrasting water availabilities

**DOI:** 10.1101/2021.07.19.453016

**Authors:** Gabriel Mulero, Harel Bacher, Uri Kleiner, Zvi Peleg, Ittai Herrmann

**Author notes:** **Corresponding author**: Ittai Herrmann, The Robert H. Smith Institute of Plant Sciences and Genetics in Agriculture, The Robert H. Smith Faculty of Agriculture, Food and Environment, The Hebrew University of Jerusalem, P.O. Box 12, 7610001 Rehovot, Israel.

## Abstract

To meet the ever-growing global population necessities, it is needed to identify climate change-relevant plant traits to integrate into breeding programs. Developing new tools for fast and accurate estimation of chlorophyll parameters, chlorophyll a (Chl-a), chlorophyll b (Chl-b) content, and their ratio (Chl-a/b), can promote breeding programs of wheat with enhanced climate adaptively. Spectral reflectance of leaves is affected by changes in pigments concentration and can be used to estimate chlorophyll parameters. The current study identified and validated the top spectral indices known and developed new vegetation indices (VIs) for Chl-a and Chl-b content estimation and used them to non-destructively estimate Chl-a/b values and compare them to hyperspectral estimations. Three wild emmer introgression lines, with contrasting drought stress responsiveness dynamics, were selected. Well and limited irrigation irrigation regimes were applied. The wheat leaves were spectrally measured with a handheld spectrometer to acquire their reflectance at the 330 to 790 nm range. Regression models based on calculated VIs as well as all hyperspectral curves were calibrated and validated against chlorophyll extracted values. The developed VIs resulted in high accuracy of Chl-a and Chl-b estimation allowing indirect non-destructive estimation of Chl-a/b with root mean square error (RMSE) values that could fit 6 to 10 times in the range of the measured values. They also performed similarly to the hyperspectral models. Altogether, we present here a new tool for a non-destructive estimation of Chl-a/b which can serve as a basis for future breeding efforts of climate-resilience wheat as well as other crops.

## Introduction

Current and projected climate change, as express in the increasing intensity of erratic climate events across extensive regions of the planet, threaten to increase food insecurity worldwide (Myers et al. 2017). To meet the ever-growing global population demands for food, feed, fibers, and bioenergy plant-based products, a significant increase in crop-plant production is required (reviewed by Sade and Peleg 2020). Thus, there is an urgent need to develop climate change resilience crop-plants with enhancing yield and nutritional quality. A fundamental aspect of such an effort is the identification of key functional traits that can be integrated into breeding programs to complement the changing agro-systems.

Chlorophyll is an important light-absorbing photosynthetic pigment largely determining the plant’s photosynthetic capacity and as consequence its growth and development (Croft et al. 2017). It includes the chlorophyll a (Chl-a), which is the primary electron donor within the reaction center, and chlorophyll-b (Chl-b), a light-harvesting accessory pigment found in the antenna complexes of the light-harvesting complexes of photosystem II (Croce and van Amerongen 2014; Croft and Chen 2018). The ratio between chlorophyll a and b (Chl-a/b) is affected by the plant’s natural senescence processes and various environmental cues (Lenaerts et al. 2019). Under water stress, the Chl-b degrades into Chl-a, leading to a higher Chl-a/b (Guo et al. 2016). Thus, developing new tools for fast and accurate estimation of chlorophyll parameters can promote breeding programs of crop-plants with enhancing climate adaptively.

Quantification of chlorophyll content in plant leaf samples was established by integrating empirical models of spectrophotometry measurement of light transmission wavelengths based on Beer-Lambert law (Moran and Porath 1980; Wellburn 1994). While resulting in accurate chlorophyll content, they are destructive, labor-intensive, time-consuming, and do not allow to study the longitudinal and spatio-temporal dynamics over time (Gitelson et al. 2006; Gitelson et al. 2003; Lopes et al. 2017). Alternatively, an *in-vivo* non-destructive optical hand-held absorbance-based total chlorophyll (TChl) meter (such as SPAD-502; Minolta corporation Ltd., Osaka, Japan) can be used. It can obtain a quick prediction of TChl content, but in many cases, the SPAD values are not calibrated to actual TChl content (Yamamoto et al. 2002; Joynson et al. 2021), and cannot estimate Chl-a, Chl-b and their ratio. Hyperspectral data is an alternative non-destructive approach, based on extracting reflectance values for hundreds of narrow spectral bands (Thenkabail et al. 2013). These data can be used for building models to determine TChl, Chl-a and Chl-b content as well as Chl-a/b. Spectral reflectance of leaves is strongly affected by changes in pigments concentration that are negatively related to spectral reflectance (Chappelle et al. 1992; Gitelson et al. 2005; Yu et al. 2015). A vegetation index (VI) is a mathematical manipulation based on reflectance values from two or more spectral bands (Bannari et al. 1995) used to estimate plant traits and monitor their health and condition (Giovos et al. 2021). Spectral data and VIs are analyzed to estimate chlorophyll content in vegetation (Bannari et al. 2007; Banerjee et al. 2020; Chen et al. 2020; Croft and Chen 2018; Gitelson 2011; Main et al. 2011; Sonobe et al. 2020).

Wheat (*Triticum* sp.) is one of the world’s most consumed crops, with production estimated at ^~^770 million tons per annum (http://www.fao.org/faostat). To meet the rising demand of the projected 9.7 billion people by 2050, an increase of at least 60% in wheat production is required (Ray et al. 2013). Wheat domestication and subsequent evolution under domestication involved a suite of complex genetic, morphological, anatomical, and physiological modifications (Abbo et al. 2014; Golan et al. 2018). Wild emmer wheat [T. *turgidum* ssp. *dicoccoides* (Körn.) Thell.], the direct allotetraploid (2n=4x=28; genome BBAA) progenitor of domesticated wheats, thrives across wide eco-geographic amplitude across the Fertile Crescent and offers ample allelic repertoire for agronomically important traits, including drought tolerance (e.g., Bacher et al. 2021; Peleg et al. 2005).

Recently we evaluated a large set of wild emmer wheat introgression lines (ILs) under contrasting water availabilities and identified promising drought tolerance strategies (Bacher et al. 2021). Here we applied a field-based evaluation of selected ILs with divergent water-stress responsiveness and tested their Chl-a/b ratio alteration in response to water stress. Our working hypothesis was that Chl-a/b can be assessed non-destructively *in-vivo* wheat leaves based on Chl-a and Chl-b contents. The specific objectives of the current study were to: ***i***) identify and validate the best VIs for Chl-a and Chl-b, ***ii***) use the best VIs to assess Chl-a/b under contrasting water availabilities, and ***iii***) compare VIs- and hyperspectral-based Chl-a/b estimation models. Altogether, we showed here for the first time, to the best of our knowledge, a new tool, based on a spectral assessment of Chl-a and Chl-b for non-destructive estimation of Chl-a/b which can serve as a basis for future breeding efforts of climate-resilience wheat, as well as other crops.

## Material and Methods

### 2.1. Plant material and experimental design

Previously we characterized a set of adaptive wild emmer introgression lines (ILs; BC_3_F_5_) for their drought responsiveness strategies. For the current study, we selected three lines (IL46, IL82, IL105) with contrasting drought stress responsiveness dynamics (Bacher et al. 2021). IL46 was characterized as highly productive and stable and exhibited high growth and gas exchange under water stress. IL82 was characterized as highly productive and exhibited high growth under water stress with phenotypic plasticity (i.e., its physiological and morphological parameters were changing due to the water stress). IL105 was characterized to have lower biomass productivity under water stress and define as high phenotypic plasticity. Each line consists of few introgressions from the wild emmer line Zavitan, with IL46 consists of six introgressions that cover 5.2% of the genome, IL82 consists of fifteen introgressions that cover 13.37% of the genome, and IL105 consist of six introgressions that cover 6.07% of the genome (Supplementary Table S1).

The plants were grown during the winter of 2019–2020 at the experimental farm of The Hebrew University of Jerusalem in Rehovot, Israel (34 47′ N, 31 54′ E: 54 m above sea level) in a plastic-covered net house. The soil is brown-red degrading sandy soil (Rhodoxeralf) composed of 76% sand, 8% silt, and 16% clay. A split-plot factorial (genotype x irrigation regime) design was employed with two irrigation regimes split into 12 sub-plots, with four replicates (total 24 plots). Each plot of 150 cm long consisted of four planted rows. Plants were spaced 10 cm, within and between rows resulting in 60 plants per plot. Two irrigation regimes were applied via a drip irrigation system: well-watered control (WW) and water-limited (WL). The WW treatment was irrigated weekly with a total amount of ^~^750 mm, whereas the WL treatment was irrigated every other week with a total amount of ^~^250mm (Supplementary Fig. S1). Water was applied during the winter months (Jan–Mar) to mimic the natural pattern of rainfall in the eastern Mediterranean region. The experiment was treated with fungicides and pesticides to control fungal pathogens or insect pests and was weeded manually once a week.

### Data collection

Spectral data were acquired by PolyPen RP410 UVIS (PSI Ltd., Drasov, Czech Republic) in contact with the adaxial leaf side. The PolyPen is a leaf contact active spectrometer covering the range of 330 to 790 nm in 1 nm intervals (8 nm band width at full width half max). Before spectral data collection and in-between measurements, white reference measurement was acquired using a spectralon panel (PSI Ltd., Drasov, Czech Republic). Each selected leaf was *in vivo* spectrally measured five times to result in an average spectrum. The youngest fully developed leaf was selected for spectral data collection and sampling, starting the third until the ninth measuring date the flag-leaf was measured. In each plot, three leaves from different plants were marked and spectrally measured to represent the plot. The spectral data acquisition was followed by leaf sample collection. The leaves were cut from the plant into an air-tight polyethylene sealed bag and then placed into an ice-filled container for up to 2 hours before further laboratory measurements.

Five leaf discs (0.8 cm diameter) were taken from each leaf and placed in a glass container with 10 ml of organic solvent (N.N Dimethylformamide) and transported into a dark 4°C incubator for 48hrs. Then, the samples were pipette into 3 ml quartz cuvettes in the UV/VIS Spectrophotometer (ST-VS-723; Lab-Fac instrument Ltd., Kowloon, Hong Kong) to acquire transmittance in two wavelengths: 647 and 664 nm. The transmittance values were used to calculate the Chl-a, Chl-b, and TChl content (mg cm^−2^), as described previously (Moran 1982). Three plants per plot were sampled, the spectral and chlorophyll contents were averaged per plot.

### Data preprocessing and analyses

Analysis of Variance (ANOVA) was used to assess the possible effects of genotype, irrigation regimes, data collection dates, and the different levels of interactions on the chlorophyll responses.

Thirty-three well-known VIs were pre-programmed to be calculated by the PolyPen sensor (Supplementary Table S2). The quality of the correlation between each of the VIs and Chl-a, Chl-b, and TChl content was evaluated by the correlation coefficient (R). VIs resulting in R absolute values equal to or higher than 0.5 were selected for linear regression to produce a coefficient of determination (R^2^) and root mean square error of prediction (RMSEP). Data sets were randomly selected with an even distribution on genotype by irrigation regime by date for all parameters based on the number (*n*) of samples involved; Chl-a (*n*=168), Chl-b (*n*=167) and TChl (*n*=166), at a 70% and 30%, calibration and validation, respectively. The calibration data sets were used to perform a linear regression analysis between each of Chl-a, Chl-b, and TChl content (dependent) and each of the selected VIs (independent) variables and determine the slope and intercept to be tested by the validation data sets. The predicted Chl-a, Chl-b, and TChl content values were compared to the observed values to obtain the calibration and validation R^2^ (Cal and Val R^2^, respectively) as well as RMSEP of calibration and validation (RMSEPC and RMSEPV, respectively) as calculated by (Herrmann et al. 2010). The % RMSEPC and % RMSEPV were calculated based on the RMSEPC and RMSEPV values out of the range of Cal and Val samples, respectively. The statistical analyses were applied in JMP 15 pro version statistical package (SAS Institute, Cary, NC, USA).

To find the best two bands combination for Chl-a, Chl-b, and TChl content spectral estimation, the normalized difference spectral index (NDSI; Inoue et al. 2008) was calculated, analyzed and ranked based on R^2^ values of a linear regression between each Chl-a, Chl-b, and TChl content (Supplementary Table S2). The highly ranked NDSI for each of the three chlorophyll parameters (i.e., Chl-a, Chl-b, and TChl content) was used for calibration and validation process as done with all selected VIs. The models’ quality was assessed by R^2^, RMSE and % RMSE. NDSI analysis was performed in R (version 3.4.1.) environment and statistical analyses were applied in JMP 14 pro version statistical package (SAS Institute, Cary, NC, USA).

In spectral datasets, as in the current study, there is high collinearity among variables (i.e., adjacent wavelengths), partial least squares regression (PLSR) is a commonly applied method (Singh et al. 2015; Wold et al. 2001; Yang et al. 2017) that used the information at all wavelengths to provide quantitative determination plant traits. To calculate the estimated trait by the regression equation, each wavelength receives a coefficient, the absolute value of these coefficients indicates the importance of each wavelength to the model. The Cal and Val samples distribution were the same as for the VIs analysis and the models’ quality was assessed by R^2^, RMSE and % RMSE. The statistical analysis was performed in python 3.8 with pandas (McKinney 2010), version 1.3, SciPy (Virtanen et al. 2020),version 1.6, and scikit-learn (Pedregosa et al. 2011), version 0.24, PLSR with NIPALS algorithm.

## Result and Discussion

To test the effect of water availability on the accuracy of various non-destructive spectral models to estimate the Chl-a/b ratio we used three selected wild emmer wheat introgression lines (ILs) that represent contrasting stress responsiveness strategies (L46, IL82 and IL105; see Bacher et al. 2021). Characterization of these ILs under two contrasting water availabilities showed a clear effect of the irrigation regime on plant height and productivity (Supplementary Table S3 and Supplementary Fig. S2). In general, under water stress, all genotypes exhibited a significant reduction in height, vegetative dry weight and the final grain yield. These results indicate that the applied water stress had a significant impact on crop production and thus can serve as a good experimental platform for studying the potential of various VIs (Supplementary Table S2) to estimate chlorophyll content.

The genotypes had a significant effect on leaf Chl-a, Chl-b, TChl, and Chl-a/b (*P*<0.0001), and the irrigation regimes affected significantly only the Chl-b and Chl-a/b (*P*=0.024 and *P*=0.021, respectively), but not on Chl-a and TChl (*P*=0.229 and *P*=0.610, respectively) (Supplementary Table S4). In wheat, Ashraf and Harris (2013) showed that Chl-a increased in response to drought, and tolerant cultivars exhibited a slight increase in the Chl-a/b. In the current study, Chl-a/b values decline along with the development of the experiment and showed non-significant differences between irrigation treatments (Supplementary Fig S3a). The spectral reflectance curves of the fully developed leaf were similar to those of the flag-leaf but in a closer look, the earlier is not following the chronological trend of the latter in the visible spectral region (Supplementary Fig. S3b). The chronological trend is visible around 730 nm. This mismatch in chronology led to a model based on the flag-leaf data alone. Leaf structure and pigment content affect leaf spectral reflectance (Gausman and Allen 1973). The relatively high mesophyll cell number per leaf area of flag-leaf in comparison to the fully developed leaf (Pyke et al. 1990) and variability in leaf thickness (Guru et al. 2017) may explain the lower reflectance of the fully developed leaves in the range of 740-780 nm. Thus, it was expected to observe a reduction in the ability of spectral data to explain the variability in Chl-a and Chl-b content while analyzing the two leaf types together. Chl-b had the biggest advantage, in R values, for analyzing only the flag-leaf spectral data rather than the two leaf types together (Supplementary Fig. S4). To improve the quality of chlorophyll spectral estimation in *in-vivo* wheat leaves and develop a standardized data collection methodology the flag-leaf data alone was further analyzed. The focus of the current study is at Chl-a/b based on Chl-a and Chl-b spectral estimation, nevertheless, TChl that is commonly assessed, was discussed as an additional relevant output. Although leaf reflectance in the visible spectral region is negatively related to chlorophyll content (Gausman and Allen 1973) the calculated VIs can be either positively or negatively related to chlorophyll content based on the bands used and the structure of the VI equation. It was hypothesized that the combination of GM1 (Gitelson and Merzlyak 1997) positively related to chlorophyll content and Carter 1 (Carter 1994) negatively related to chlorophyll content will improve the chlorophyll estimation quality. Two VIs were developed (#38 and #39 in Supplementary Table S2) and proved to be at the top of TChl estimation and with higher R^2^ and smaller RMSE than each of their components alone (Table 1).

**Table 1:**
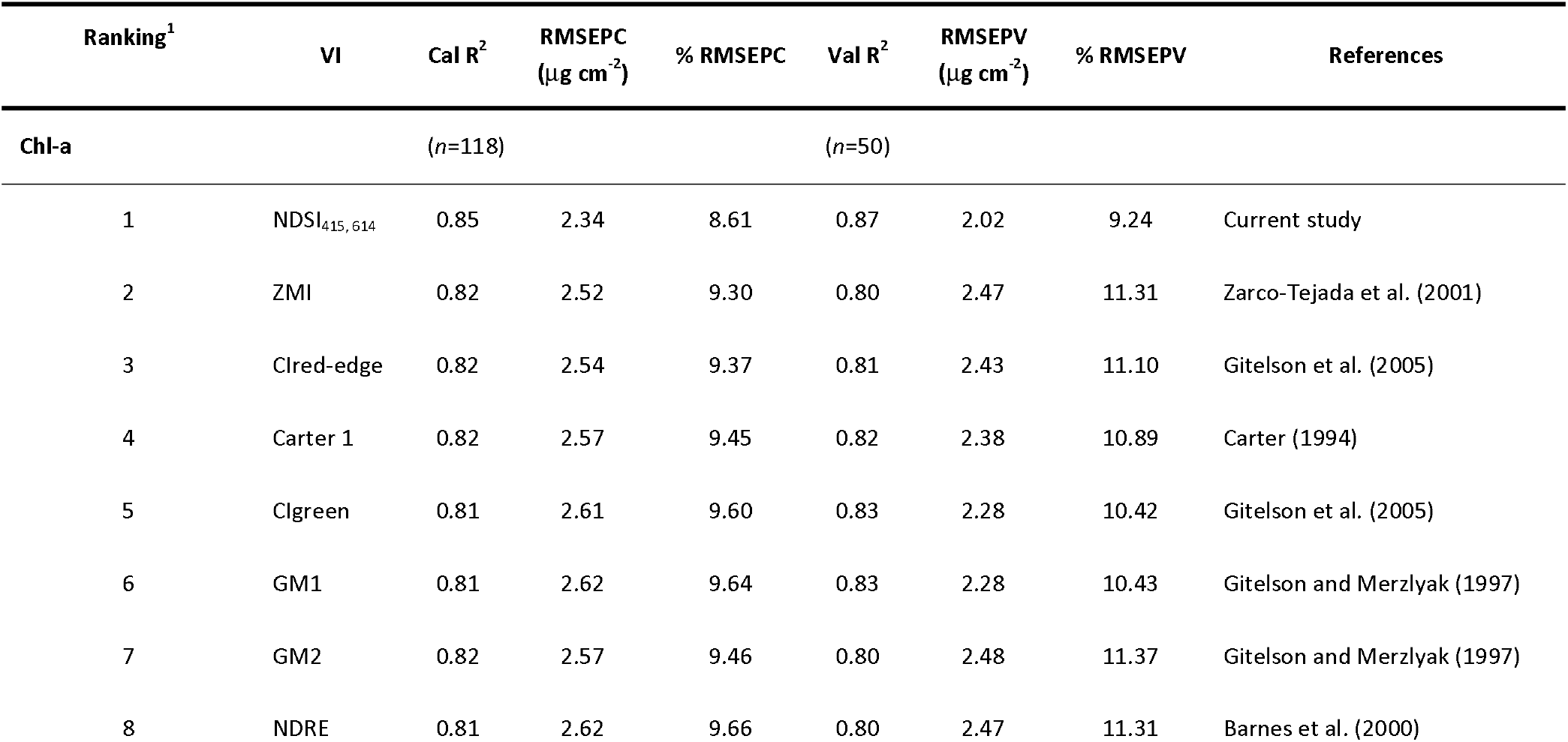
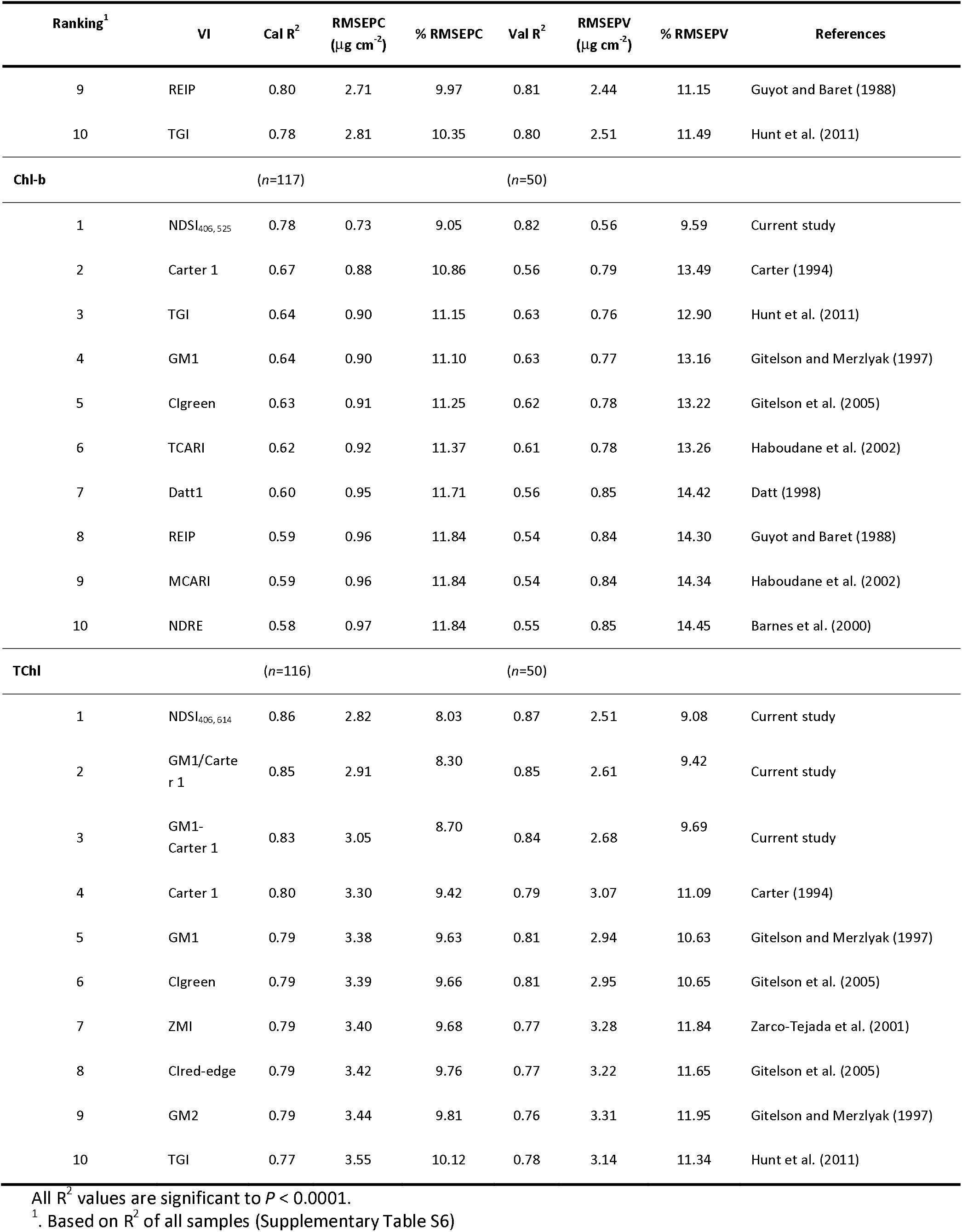
The top ten predictions for Chl-a, Chl-b, and TChl are presented based on all 33 VIs, NDSI generated VIs from the leaf reflectance spectra by the PolyPen sensor.

## Vegetation indices (VIs) enable non-destructive and accurate assessment of leaf chlorophyll content

The R^2^ distribution of the two bands combinations to assess Chl-a, Chl-b and TChl content are presented in heat maps (Fig. 1a, c and e). As expected the heat maps are similar for the three chlorophyll parameters (Datt 1998) but with smaller areas with relatively high R^2^ values for Chl-b. The selected bands for Chl-a and TChl are very similar as can be expected since Chl-a is the major parameter in TChl. It is important to mention that bands adjacent to the selected ones will also show high R^2^ values (Table S5). The best two bands combination for the Chl-b estimation was 406 and 525 nm, supported by Ustin and Jacquemoud (2020) reporting Chl-b absorption peaks in the blue and green regions. Analyzing the averaged spectral and destructive replicates per day (Fig. 1 b, d and f) resulted in bigger R^2^ values than the non-averaged ones (Fig. 1a, c and e). All VIs were correlated to Chl-a, Chl-b and TChl and ranked based on R^2^ values (Supplementary Table S6). The top 10 performing VIs (Supplementary Table S6) used for Chl-a, Chl-b, and TChl estimation resulted in similar Cal and Val R^2^ as well as RMSE values (Table 1). The RMSE values (Fig. 3 and Table 1) of indices developed in the current study are smaller than 10% of the active ranges of Chl-a, TChl and even of Chl-b. Previous studies as detailed by Hallik et al. (2017) presented smaller R^2^ and bigger RMSE values of Chl-a and Chl-b estimation based on spectral data and stated that studies estimating Chl-b used VIs that were more strongly correlated to Chl-a. In the current study, VIs were developed specifically for Chl-a as well as Chl-b estimation and are at the top in their category (Table 1). Banerjee et al. (2020) developed a VI highly correlating with Chl-a, Chl-b and TChl concentrations based on wheat canopy side view imagery, acquired in a semi-controlled indoors environment. This VI used spectral regions related to nitrogen (1654 nm) and chlorophyll (727 nm). Sonobe et al. (2020) applied spectral methods to estimate leaf Chl-a and Chl-b in wasabi grown in a semi-controlled environment indoors and resulted in RMSE values similar to the values achieved in the current study (Table 1). The current study presented improvement (in terms of R^2^ and RMSE) in the ability to estimate leaf Chl-a, Chl-b and TChl content values in wheat grown under field conditions and used the spectrally estimated values to calculate Chl-a/b (Fig. 2). Spectral estimation of Chl-a/b was rarely published, Sonobe et al. (2020) directly estimated Chl-a/b by spectral means resulting in RMSE values ranging from 0.13 to 0.6. In the current study, the RMSE can fit five to six times in the range of measured Chl-a/b values (Fig. 2a). To the best of our knowledge, the approach of spectrally estimating Chl-a and Chl-b values in *in-vivo* wheat leaves to calculate Chl-a/b was not published yet. The VIs developed in the current study were using two or few spectral bands while the PLSR applied hundreds of spectral bands to estimate Chl-a and Chl-b values.

**Figure 1.**
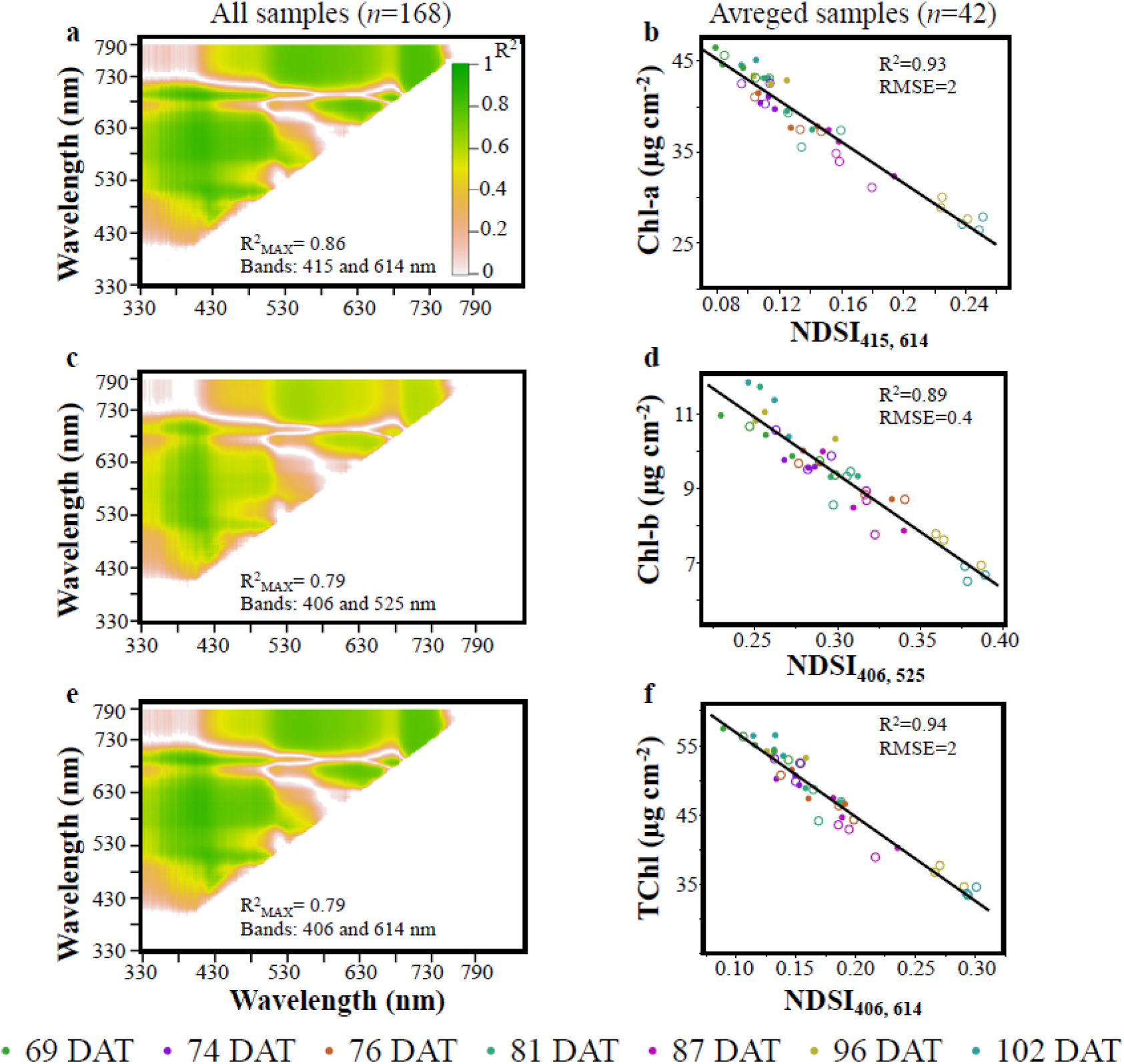
All possible two-band combinations by normalized difference vegetation index (NDSI) heat maps R^2^ values (*P*<0.0001) for regression with Chl-a, Chl-b, and TChl content on all 168 samples (**a, c, and e**). Linear regression of Chl-a, Chl-b, and TChl (*n*=42) with their corresponding highest-ranking NDSI based on averaged NDSI values per genotype, irrigation regime and DAT (**b, d, and f**) the black lines are the trend line. Chl-a (**a-b**), Chl-b (**c-d**), and TChl (**e-f**). Hollow and filled dots stand for water-limited (WL) and well-watered (WW), respectively. RMSE stands for root mean square error; DAT stands for days after transplant.

**Figure 2.**
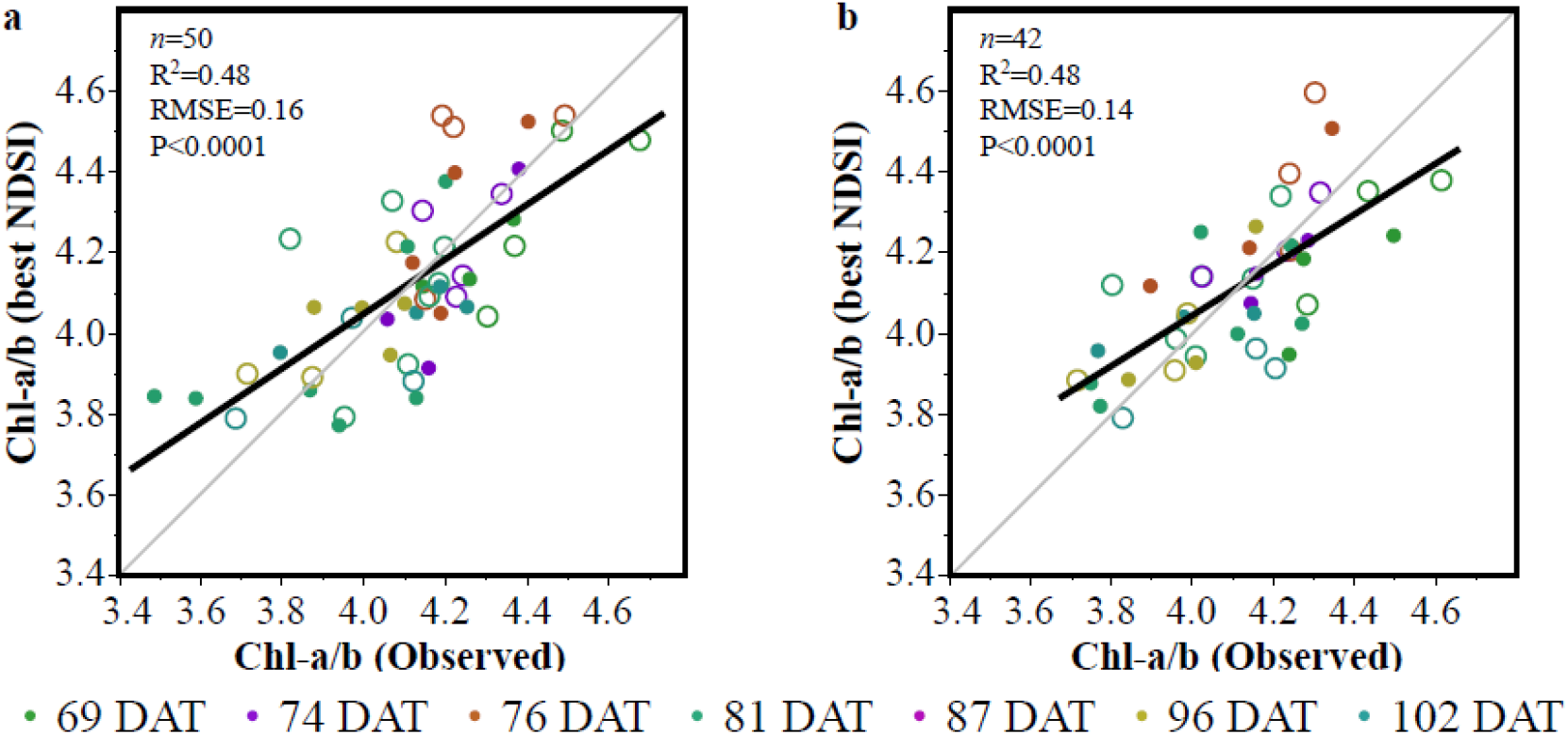
Quality of Chl-a/b spectral estimation by normalized difference spectral index (NDSI). The Chl-a/b (observed) data was acquired by extraction of Chl-a and Chl-b. The Chl-a/b (best NDSI) data was calculated by applying the top (NDSI estimating Chl-a and Chl-b (Table 1). The Val data set is presented in Table 1 (**a**). Averaged values per genotype, irrigation regime and DAT as presented in Fig. 1 b, d, and f (**b**). DAT stands for days after transplant. Hollow and filled dots stand for water-limited (WL) and well-watered (WW), respectively; the black lines are the trend lines and the gray lines are the 1:1 lines.

**Figure 3.**
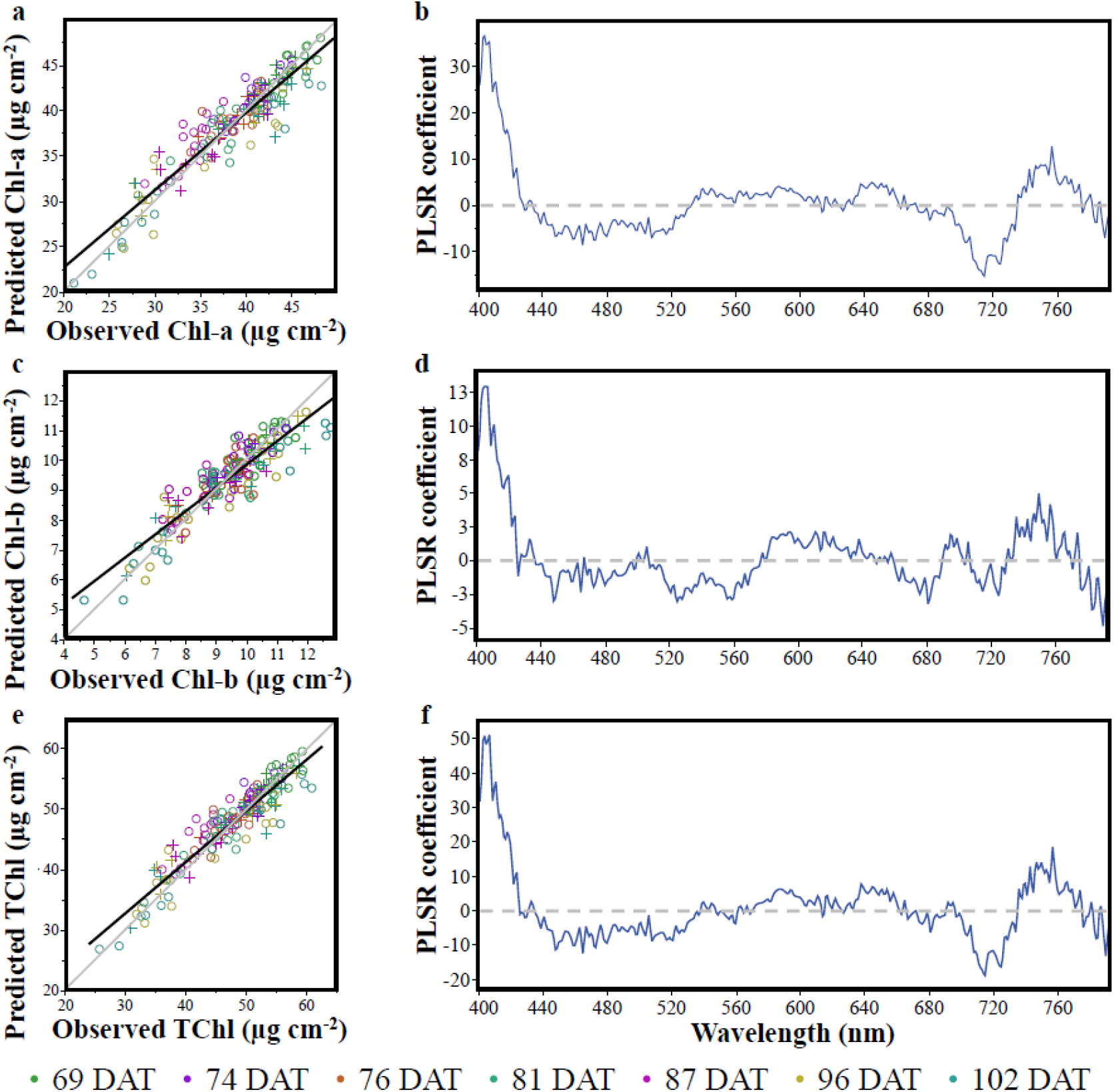
Chl-a, Chl-b, and TChl content estimation by partial least squares (PLSR) models. Observed vs predicted chlorophyll values (**a, c, and e)**, solid black lines are calibration (Cal) best fit lines and 1:1 lines are gray, Circle and plus markers represent Cal and Val samples, respectively. The PLSR coefficients for each of the models (**b, d, and f**), solid blue lines are coefficient values and dashed gray lines are the zero-coefficient value. Chl-a (**a-b**), Chl-b (**c-d**), and TChl (**e-f**).

## The partial least squares regression (PLSR) model did not have an advantage on the vegetation indices (VIs)

The RMSE obtained for the estimated Chl-a, Chl-b and TChl concentrations by PLSR models (Table 2) are small enough to fit 12, 14, and 11 times, respectively, in the observed range of concentrations (Fig. 3 a, c, and e). The estimation quality by PLSR and VIs (Table 1) in terms of R^2^ and RMSE was similar (Herrmann et al. 2011) for each of the chlorophylls. The VIs based on two to a few spectral bands and the PSLR based on 391 spectral bands resulted similarly showing no advantage to either of them. The PLSR coefficients of the models (Fig. 3 b, d, and f) are all showing the importance of the shortest wavelengths in agreement with the 406 and 415 nm wavelengths selected by the NDSIs (Fig. 1 a, c, and e). The combined effect of the Chl-a and Chl-b coefficients can be seen in the TChl coefficients in the enhancement in the shortest wavelengths as well as the contradicting trends between 500 and 550 nm as well as around 700 nm. These insights support the stability of the models to be able to assess Chl-a and Chl-b and their sum as well as the ability to obtain Chl-a/b estimation (Fig. 4). The Chl-a/b estimation based on the PLSR models (Fig. 4) is showing improved R^2^ and RMSE in comparison to the VIs for the Val data set (Fig. 2a) as well as for the averaged samples (Fig. 2b) as was done for the VIs (Fig. 1 b, d, and f). As expected the averaged samples data resulted in the smallest RMSE value. The NDSI results (Fig. 1 a, c, and e) support using the spectral range of 400 to 790 nm for the PLSR analysis.

**Table 2.** The Chl-a, Chl-b, and TChl estimation based on PLSR model.

| Chlorophyll | Cal $R^2$ | RMSEPC<br>($\mu\text{g cm}^{-2}$) | % RMSEPC | Val $R^2$ | RMSEPV<br>($\mu\text{g cm}^{-2}$) | % RMSEPV |
| --- | --- | --- | --- | --- | --- | --- |
| Chl-a | 0.88 | 2.11 | 7.76 | 0.86 | 2.08 | 9.52 |
| Chl-b | 0.80 | 0.66 | 8.10 | 0.81 | 0.57 | 9.63 |
| TChl | 0.87 | 2.68 | 7.63 | 0.86 | 2.51 | 9.05 |

**Figure 4.**
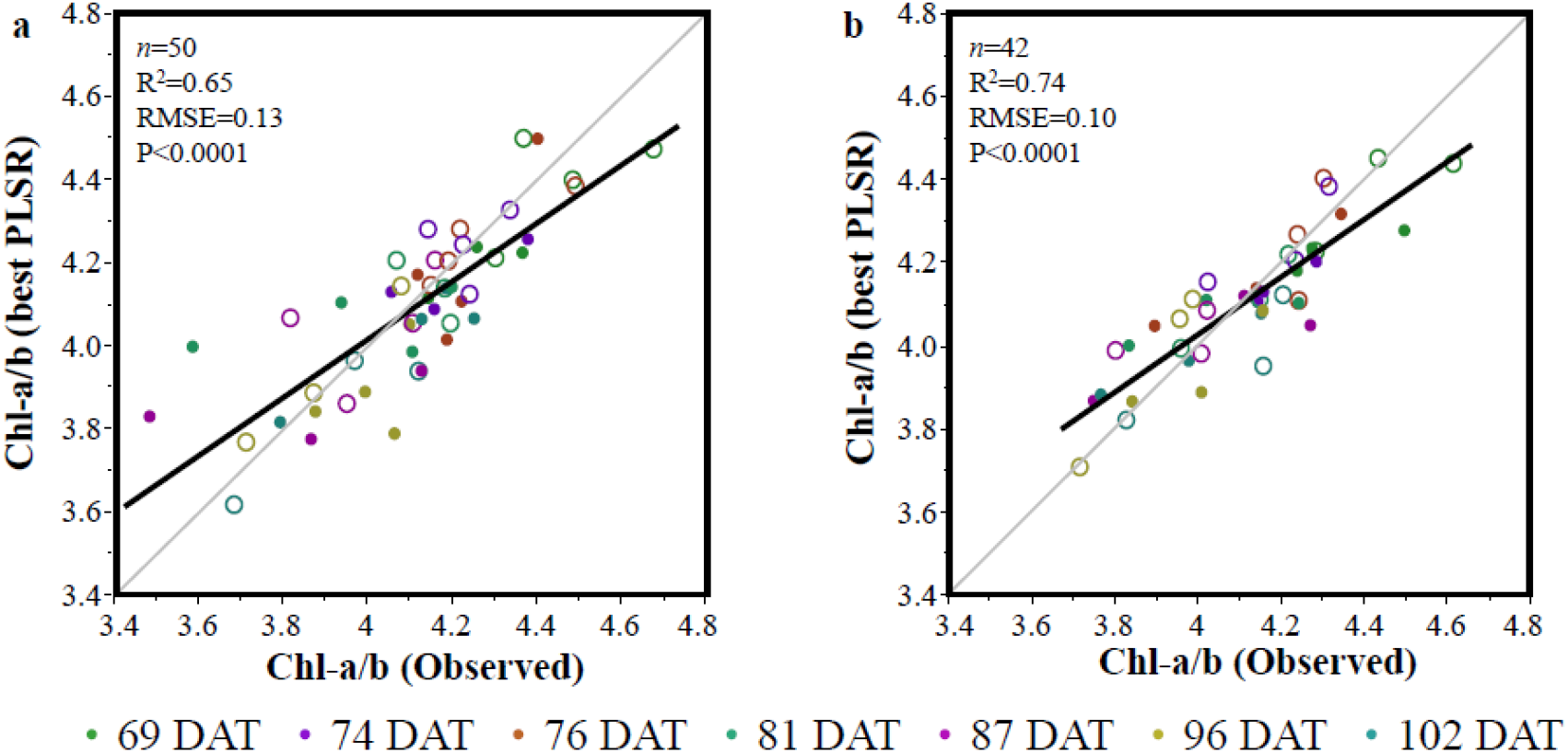
Quality of Chl-a/b spectral estimation. The Chl-a/b (observed) data was acquired by extraction of Chl-a and Chl-b. The Chl-a/b (best PLSR) data was calculated by applying the PLSR models estimating Chl-a and Chl-b (Table 2). The Val data set is presented in Table 2. Hollow and filled dots stand for water-limited (WL) and well-watered (WW), respectively (**a**). Averaged values per genotype, irrigation regime and DAT as presented in Fig. 3 b, d, and f (**b**). The black lines are the trend lines and the gray lines are the 1:1 lines. Hollow and filled dots stand for water-limited (WL) and well-watered (WW), respectively.

## Concluding remarks

The current study aimed to spectrally assess Chl-a and Chl-b to identify the best VI or VIs for indirectly estimate Chl-a/b. While there were no significant differences in Chl-a/b values between WW and WL treatments, the active range of the measured or predicted Chl-a/b values were five to six times the RMSE values for the non-averaged samples. Thus, it was concluded that: ***i***) the new VIs that were developed in the current study resulted in high accuracy of Chl-a and Chl-b estimation; ***ii***) the developed VIs were able to indirectly estimate Chl-a/b; and ***iii***) the VIs developed in the current study performed similarly to the PLSR. The model quality achieved by VIs developed in the current study in comparison to PLSR supports sensors with few spectral bands for practical use while hyperspectral sensors are used for research to identify these spectral bands. The developed models should be tested in additional crops for breeding projects. Under more severe water stress scenarios, resulting in a wider range of Chl-a/b values, the models are expected to perform even better than in the current study. This concept of Chl-a and Chl-b direct estimation to indirectly assess Chl-a/b should be tested also for canopy level spectral data collection. The current study presented a new tool, based on a spectral assessment of Chl-a and Chl-b for non-destructive estimation of Chl-a/b which can serve as a basis for future wheat breeding efforts. The ability to assess Chl-a/b *in-vivo*, non-destructively by spectral sensing will improve breeding projects efficiency.

## Supporting information

Sup. tabs and figs

## Abbreviations

ANOVA: Analysis of variance
Cal: Calibration
Chl-a: Chlorophyll a
Chl-a/b: Chlorophyll a and chlorophyll b ratio
Chl-b: Chlorophyll b
DAT: Days after transplant
ILs: Introgression lines
*n*: number of samples
NDSI: Normalized difference spectral index
PLSR: Partial least squares regression
R: Correlation coefficient
R^2^: Coefficient of determination
RMSE: Root means square error
RMSEP: Root mean square error of prediction
RMSEPC: Root means square error of prediction of calibration
RMSEPV: Root means square error of prediction of validation
TChl: Total chlorophyll
Vai: Validation
Vis: Vegetation indices
WL: Water-limited
WW: Well-watered

## Acknowledgments

This study was partially supported by the U.S. Agency for International Development Middle East Research and Cooperation (grant # M34-037), and the Dutch Ministry of Foreign Affairs under Dutch development / foreign policy (Project WheatMAX), the Hebrew University of Jerusalem Intramural Research Fund Career Development, and the International School of Agricultural Sciences.

## Author contributions statement

G.M., H.B., Z.P. and I.H. designed the study. G.M., H.B. and U.K. performed the experiments and analyzed the data. G.M., H.B., Z.P. and I.H. wrote the manuscript. All authors discussed the results and their implications and revised the manuscript.

