## Supplementary material for "Spectral estimation of *in-vivo* wheat chlorophyll a/b ratio under contrasting water availabilities": Sup. tabs and figs

**Supplementary data**

**Table S1.** List of wild emmer introgression lines and their genomic composition. Description of the the Wild Introgression lines physical position annotated to the Zavitan genome

| **Introgression line** | **Chr** | **Introgression Stat Position (bp)** | **Introgression End Position (bp)** | **Introgression Size** |
| --- | --- | --- | --- | --- |
| 46 | 3A | 33195099 | 49729314 | 16534215 |
| 46 | 3A | 64995502 | 507936763 | 442941261 |
| 46 | 3B | 744946081 | 792129698 | 47183617 |
| 46 | 4B | 463426 | 9079844 | 8616418 |
| 46 | 5A | 543208564 | 548003918 | 4795354 |
| 46 | 7B | 631850491 | 684067060 | 52216569 |
| 82 | 1B | 609661262 | 647845285 | 38184023 |
| 82 | 2A | 73074586 | 662241021 | 589166435 |
| 82 | 2B | 4002969 | 21294555 | 17291586 |
| 82 | 2B | 80709457 | 392300499 | 311591042 |
| 82 | 2B | 603378597 | 631625352 | 28246755 |
| 82 | 3A | 5653 | 108315443 | 108309790 |
| 82 | 3A | 438089358 | 596670337 | 158580979 |
| 82 | 3B | 4676652 | 23716825 | 19040173 |
| 82 | 4A | 604094595 | 629409087 | 25314492 |
| 82 | 5A | 11194 | 6677852 | 6666658 |
| 82 | 5A | 520160421 | 584510015 | 64349594 |
| 82 | 5A | 656443879 | 698099510 | 41655631 |
| 82 | 5B | 686587813 | 711812306 | 25224493 |
| 82 | 6B | 118323794 | 155690123 | 37366329 |
| 82 | 7B | 485326 | 659068 | 173742 |
| 105 | 1A | 47666533 | 593106089 | 545439556 |
| 105 | 3A | 643724679 | 698744469 | 55019790 |
| 105 | 4A | 703447171 | 710456757 | 7009586 |
| 105 | 6A | 568983599 | 582391045 | 13407446 |
| 105 | 7A | 1132235 | 14374334 | 13242099 |
| 105 | 7A | 90855345 | 124467146 | 33611801 |

**Table S2*.*** Vegetation indices (VIs) that were adopted from relevant literature and pre-programmed in the sensors, normalized difference spectral index (NDSI) and VIs developed in the current study.

| **#** | **Vegetation Index** | **Formula** | **References** |
| --- | --- | --- | --- |
| 1 | NDVI | (R_780_ – R_630_)/(R_780_ + R_630_) | Rouse et al., (1974) |
| 2 | SR | R_780_/R_630_ | Jordan, (1969) |
| 3 | MCARI 1 | 1.2*[2.5*(R_780_ – R_670_) – 1.3*(R_780_ – R_550_)] | Haboudane et al., (2002) |
| 4 | OSAVI | (1+0.16)*[(R_780_ – R_670_)/( R_780_ + R_670_ +0.16)] | Rondeaux et al., (1996) |
| 5 | G | R_554_/R_677_ | Smith et al., (1995) |
| 6 | MCARI | [(R_700_ – R_670_) – 0.2*(R_700_ – R_550_)]*(R_700_/R_670_) | Daughtry et al., (2000) |
| 7 | TCARI | 3*[(R_700_ – R_670_) – 0.2*(R_700_ – R_550_)*(R_700_/R_670_)] | Haboudane et al., (2002) |
| 8 | TVI | 0.5*[120*(R_700_ – R_550_) – 200*( R_670_ – R_550_)] | Broge & Leblanc, (2001) |
| 9 | ZMI | R_750_/R_710_ | Zarco-Tejada et al., (2001) |
| 10 | SRPI | R_430_/R_680_ | Peñuelas & Filella, (1995) |
| 11 | NPQI | (R_415_ – R_435_)/(R_415_ + R_435_) | Barnes et al., (1992) |
| 12 | PRI | (R_531_ – R_570_)/(R_531_ + R_570_) | Gamon et al., (1992) |
| 13 | NPCI | (R_680_ – R_430_)/(R_680_ + R_430_) | Peñuelas et al., (1994) |
| 14 | Carter 1 | R_695_/R_420_ | Carter, (1994) |
| 15 | Carter 2 | R_695_/R_760_ | Carter, (1994) |
| 16 | Lic 1 | (R_780_ – R_680_)/(R_780_ + R_680_) | Lichtenthaler, (1996) |
| 17 | Lic 2 | R_440_/R_690_ | Lichtenthaler, (1996) |
| 18 | SIPI | (R_780_ – R_450_)/(R_780_ + R_450_) | Penuelas et al., (1995) |
| 19 | GM 1 | R_750_/R_550_ | Gitelson & Merzlyak, (1997) |
| 20 | GM 2 | R_750_/R_700_ | Gitelson & Merzlyak, (1997) |
| 21 | ARI 1 | 1/R_500_ – 1/R_700_ | Gitelson et al., (2001) |
| 22 | ARI 2 | R_790_*(1/R_550_ – 1/R_700_) | Gitelson et al., (2001) |
| 23 | CRI 1 | 1/R_510_ – 1/R_550_ | Gitelson et al., (2002) |
| 24 | CRI 2 | 1/R_510_ – 1/R_700_ | Gitelson et al., (2002) |
| 25 | RDVI | (R_780_ – R_670_)/(R_780_ + R_670_)^0.5^ | Roujean & Breon, (1995) |
| 26 | CLred-edge | (R_783_/R_705_)-1 | Gitelson et al., (2005) |
| 27 | CLgreen | (R_783_/R_560_)-1 | Gitelson et al., (2005) |
| 28 | REIP | 700 + 40*[((R_670_ – R_780_)/2) – R_700_)/(R_740_ – R_700_)] | Guyot et al., (1988) |
| 29 | NDRE | (R_790_ – R_720_)/(R_790_ + R_720_) | Barnes et al.,(2000) |
| 30 | TGI | -0.5*[190*(R_670_ – R_550_) – 120*(R_670_ – R_480_)] | Raymond Hunt et al., (2011) |
| 31 | Datt1 | R_672_/(R_550_*R_708_) | Datt, (1998) |
| 32 | Datt2 | R_672_/R_550_ | Datt, (1998) |
| 33 | PSRI | R_672_/(R_550_ + R_708_) | Merzlyak et al., (1999) |
| 34 | NDSI | (R_i_ – R_j_)/(R_i_ + R_j_) | Inoue et al., (2008) |
| 35 | NDSI Chl-a | (R_415_ – R_614_)/(R_415_ + R_614_) | Current study |
| 36 | NDSI Chl-b | (R_406_ – R_525_)/(R_406_ + R_525_) | Current study |
| 37 | NDSI TChl | (R_406_ – R_614_)/(R_406_ + R_614_) | Current study |
| 38 | GM1- Carter 1 | (R_750_/R_550_)–(R_695_/R_420_^)^ | Current study |
| 39 | GM1/ Carter 1 | (R_750_/R_550_)(R_695_/R_420_^)^ | Current study |

R is the reflectance in the subscripted wavelength in nm; i and j represent combinations of two separate wavelengths between 330 - 790 nm.

**References (Table S2)**

Barnes, E. M., & Clarke, T.R. and Richards, S. E. (2000). Coincident detection of crop water stress, nitrogen status and canopy density using ground-based multispectral data. *In P. C. Robert, R. H. Rust, & W. E.Larson (Eds.), Proceedings of the Fifth International Conference on Precision Agriculture*. https://naldc.nal.usda.gov/download/4190/PDF

Barnes, J. D., Balaguer, L., Manrique, E., Elvira, S., & Davison, A. W. (1992). A reappraisal of the use of DMSO for the extraction and determination of chlorophylls a and b in lichens and higher plants. *Environmental and Experimental Botany*, *32*(2), 85–100. https://doi.org/10.1016/0098-8472(92)90034-Y

Broge, N. H., & Leblanc, E. (2001). Comparing prediction power and stability of broadband and hyperspectral vegetation indices for estimation of green leaf area index and canopy chlorophyll density. *Remote Sensing of Environment*, *76*(2), 156–172. https://doi.org/10.1016/S0034-4257(00)00197-8

Carter, G. A. (1994). Ratios of leaf reflectances in narrow wavebands as indicators of plant stress. *International Journal of Remote Sensing*, *15*(3), 517–520. https://doi.org/10.1080/01431169408954109

Datt, B. (1998). Remote sensing of chlorophyll a, chlorophyll b, chlorophyll a+b, and total carotenoid content in eucalyptus leaves. *Remote Sensing of Environment*, *66*(2), 111–121. https://doi.org/10.1016/S0034-4257(98)00046-7

Daughtry, C. S. ., Walthall, C. ., M.S, K., E., B. de C., & McMurtrey, J. E. (2000). Estimating Corn Leaf Chlorophyll Concentration from Leaf and Canopy Reflectance. *Remote Sensing of Environment*, *74*(2), 229–239. https://doi.org/10.1016/S0034-4257(00)00113-9

Gamon, J. A., Penuelas, J., & Field, C. B. (1992). *A Narrow-Waveband Spectral Index That Tracks Diurnal Changes in Photosynthetic Efficiency*. *41*, 35–44. https://doi.org/10.1088/0305-4470/24/13/001

Gitelson, A. A., & Merzlyak, M. N. (1997). Remote estimation of chlorophyll content in higher plant leaves. *International Journal of Remote Sensing*, *18*(12), 2691–2697. https://doi.org/10.1080/014311697217558

Gitelson, Anatoly A., Merzlyak, M. N., & Chivkunova, O. B. (2001). Optical Properties and Nondestructive Estimation of Anthocyanin Content in Plant Leaves¶. *Photochemistry and Photobiology*, *74*(1), 38. https://doi.org/10.1562/0031-8655(2001)074<0038:opaneo>2.0.co;2

Gitelson, Anatoly A., Viña, A., Ciganda, V., Rundquist, D. C., & Arkebauer, T. J. (2005). Remote estimation of canopy chlorophyll content in crops. *Geophysical Research Letters*, *32*(8), 1–4. https://doi.org/10.1029/2005GL022688

Gitelson, Anatoly A., Zur, Y., Chivkunova, O. B., & Merzlyak, M. N. (2002). Assessing Carotenoid Content in Plant Leaves with Reflectance Spectroscopy¶. *Photochemistry and Photobiology*, *75*(3), 272. https://doi.org/10.1562/0031-8655(2002)075<0272:accipl>2.0.co;2

Guyot, G., Baret, F., & Major, D. J. (1988). High spectral resolution: Determination of spectral shifts between the red and infrared. *International Archives of Photogrammetry and Remote Sensing*, *11*(1), 750–760. https://doi.org/10.1093/mind/VII.25.101

Haboudane, D., Miller, J. R., Tremblay, N., Zarco-Tejada, P. J., & Dextraze, L. (2002). Integrated narrow-band vegetation indices for prediction of crop chlorophyll content for application to precision agriculture. *Remote Sensing of Environment*, *81*(2–3), 416–426. https://doi.org/10.1016/S0034-4257(02)00018-4

Inoue, Y., Peñuelas, J., Miyata, A., & Mano, M. (2008). Normalized difference spectral indices for estimating photosynthetic efficiency and capacity at a canopy scale derived from hyperspectral and CO2 flux measurements in rice. *Remote Sensing of Environment*, *112*(1), 156–172. https://doi.org/10.1016/j.rse.2007.04.011

Jordan, C. F. (1969). Derivation of Leaf-Area Index from Quality of Light on the Forest Floor. *Ecology*, *50*(4), 663–666. https://doi.org/10.2307/1936256

Lichtenthaler, H. K. (1996). The Stress Concept in Plants: An Introduction. *Plant Physiology and Biochemistry*, *148*, 4–14. https://doi.org/10.1111/j.1749-6632.1998.tb08993.x

Merzlyak, M. N., Gitelson, A. A., Chivkunova, O. B., & Rakitin, V. Y. (1999). Non-destructive optical detection of pigment changes during leaf senescence and fruit ripening. *Physiologia Plantarum*, *106*(1), 135–141. https://doi.org/10.1034/j.1399-3054.1999.106119.x

Penuelas, J., Baret, F., & Filella, I. (1995). Semi-empirical indices to assess carotenoids/chlorophyll a ratio from leaf spectral reflectance. *Photosynthetica*, *31*(2), 221–230.

Peñuelas, J., & Filella, I. (1995). Reflectance assessment of mite effects on apple trees. *International Journal of Remote Sensing*, *16*(14), 2727–2733. https://doi.org/10.1080/01431169508954588

Peñuelas, J., Gamon, J. A., Fredeen, A. L., Merino, J., & Field, C. B. (1994). Reflectance indices associated with physiological changes in nitrogen- and water-limited sunflower leaves. *Remote Sensing of Environment*, *48*(2), 135–146. https://doi.org/10.1016/0034-4257(94)90136-8

Raymond Hunt, E., Daughtry, C. S. T., Eitel, J. U. H., & Long, D. S. (2011). Remote sensing leaf chlorophyll content using a visible band index. *Agronomy Journal*, *103*(4), 1090–1099. https://doi.org/10.2134/agronj2010.0395

Rondeaux, G., Steven, M., & Baret, F. (1996). Optimization of soil-adjusted vegetation indices. *Remote Sensing of Environment*, *55*(2), 95–107. https://doi.org/10.1016/0034-4257(95)00186-7

Roujean, J. L., & Breon, F. M. (1995). Estimating PAR absorbed by vegetation from bidirectional reflectance measurements. *Remote Sensing of Environment*, *51*(3), 375–384. https://doi.org/10.1016/0034-4257(94)00114-3

Rouse, J. W., Haas, R. H., Schell, J. A., Deering, D. W., & Harlan, J. C. (1974). *The, Monitoring Advancement, Vernal Vegetation, O F Natural*. *September 1972*.

Smith, R., Adams, J., Stephens, D., & Hick, P. (1995). Forecasting wheat yield in a Mediterranean-type environment from the NOAA satellite. *Crop \& Pasture Science*, *46*, 113–125.

Zarco-Tejada, P. J., Miller, J. R., Noland, T. L., Mohammed, G. H., & Sampson, P. H. (2001). Scaling-up and model inversion methods with narrowband optical indices for chlorophyll content estimation in closed forest canopies with hyperspectral data. *IEEE Transactions on Geoscience and Remote Sensing*, *39*(7), 1491–1507. https://doi.org/10.1109/36.934080

**Table S3**. Two-way ANOVA table of main effects and interactions effect on Height, vegetative biomass (VegBM), and grain yield (GY).

| **Source** | **DF** | **Sum of Squares** | **Mean Square** | **F Ratio** | **P value (F)** |
| --- | --- | --- | --- | --- | --- |
| **Height** |  |  |  |  |  |
| G | 2 | 157.80 | 78.90 | 7.40 | 0.0045* |
| T | 1 | 475.26 | 475.26 | 44.58 | <.0001* |
| G*T | 2 | 17.59 | 8.80 | 0.83 | 0.4541 |
| **VegBM** |  |  |  |  |  |
| G | 2 | 24432.27 | 12216.1 | 1.90 | 0.1781 |
| T | 1 | 888450.3 | 888450.2 | 138.32 | <.0001* |
| G*T | 2 | 13702.04 | 6851 | 1.07 | 0.365 |
| **GY** |  |  |  |  |  |
| G | 2 | 39838.1 | 19919 | 1.51 | 0.2472 |
| T | 1 | 1138745.2 | 1138745 | 86.43 | <.0001* |
| G*T | 2 | 43264.8 | 21632 | 1.64 | 0.2213 |

G and T represents genotypes, and irrigation regimes, respectively.

**Table S4.** A Three-way ANOVA table of main effects and interactions effect on Chl-a, Chl-b, TChl and Chl-a/b.

| **Source** | **d.f.** | **Mean Square** | **F Ratio** | ***P-*value (F)** |
| --- | --- | --- | --- | --- |
| **Chl-a** |  |  |  |  |
| G | 2 | 0.000064 | 18.32 | <0.0001 |
| T | 1 | 0.000001 | 0.26 | 0.6106 |
| D | 6 | 0.000250 | 71.12 | <.0001 |
| G*T | 2 | 0.000006 | 1.59 | 0.2086 |
| G*D | 12 | 0.000009 | 2.60 | 0.004 |
| T*D | 6 | 0.000024 | 6.76 | <0.0001 |
| G*T*D | 12 | 0.000010 | 2.74 | 0.0025 |
| **Chl-b** |  |  |  |  |
| G | 2 | 0.0000143 | 48.19 | <0.0001 |
| T | 1 | 0.0000015 | 5.20 | 0.0243 |
| D | 6 | 0.0000323 | 109.23 | <0.0001 |
| G*T | 2 | 0.0000008 | 2.67 | 0.073 |
| G*D | 12 | 0.0000009 | 3.04 | 0.0009 |
| T*D | 6 | 0.0000017 | 5.73 | <0.0001 |
| G*T*D | 12 | 0.0000009 | 3.07 | 0.0008 |
| **TChl** |  |  |  |  |
| G | 2 | 0.00014 | 26.59 | <0.0001 |
| T | 1 | 0.00001 | 1.46 | 0.2294 |
| D | 6 | 0.00040 | 76.30 | <0.0001 |
| G*T | 2 | 0.00001 | 2.04 | 0.134 |
| G*D | 12 | 0.00002 | 3.42 | 0.0002 |
| T*D | 6 | 0.00004 | 7.93 | <0.0001 |
| G*T*D | 12 | 0.00001 | 2.77 | 0.0023 |
| **Chl-a/b** |  |  |  |  |
| G | 2 | 1.24 | 69.14 | <0.0001 |
| T | 1 | 0.10 | 5.47 | 0.021 |
| D | 6 | 6.12 | 341.24 | <0.0001 |
| G*T | 2 | 0.04 | 2.13 | 0.1234 |
| G*D | 12 | 0.02 | 1.36 | 0.1914 |
| T*D | 6 | 0.04 | 2.19 | 0.0487 |
| G*T*D | 12 | 0.04 | 2.39 | 0.0082 |

T, G and D represent irrigation regimes, genotypes, and dates.

**Table S5**. Top 20 pairs of bands from the NDSI analysis in relation to Chl-a, Chl-b and TChl.

|  | **Chl-a (μg cm^-2^)** | | | **Chl-b (μg cm^-2^)** | | | **TChl (μg cm^-2^)** | | |
| --- | --- | --- | --- | --- | --- | --- | --- | --- | --- |
| **Ranking** | **Band i** | **Band j** | **R^2^** | **Band i** | **Band j** | **R^2^** | **Band i** | **Band j** | **R^2^** |
| 1 | 415 | 614 | 0.86 | 406 | 525 | 0.79 | 406 | 614 | 0.86 |
| 2 | 415 | 613 | 0.85 | 406 | 524 | 0.79 | 406 | 613 | 0.85 |
| 3 | 415 | 615 | 0.85 | 406 | 521 | 0.79 | 406 | 602 | 0.85 |
| 4 | 415 | 609 | 0.85 | 406 | 520 | 0.79 | 406 | 615 | 0.85 |
| 5 | 415 | 602 | 0.85 | 406 | 522 | 0.79 | 404 | 614 | 0.85 |
| 6 | 415 | 612 | 0.85 | 406 | 526 | 0.79 | 406 | 596 | 0.85 |
| 7 | 415 | 603 | 0.85 | 404 | 525 | 0.79 | 406 | 597 | 0.85 |
| 8 | 415 | 601 | 0.85 | 406 | 523 | 0.79 | 406 | 603 | 0.85 |
| 9 | 415 | 605 | 0.85 | 406 | 519 | 0.79 | 406 | 609 | 0.85 |
| 10 | 415 | 604 | 0.85 | 406 | 528 | 0.79 | 406 | 605 | 0.85 |
| 11 | 415 | 608 | 0.85 | 404 | 524 | 0.79 | 406 | 601 | 0.85 |
| 12 | 415 | 610 | 0.85 | 406 | 527 | 0.79 | 406 | 606 | 0.85 |
| 13 | 416 | 614 | 0.85 | 406 | 569 | 0.79 | 406 | 604 | 0.85 |
| 14 | 415 | 606 | 0.85 | 406 | 518 | 0.79 | 406 | 589 | 0.85 |
| 15 | 414 | 614 | 0.85 | 406 | 568 | 0.79 | 406 | 608 | 0.85 |
| 16 | 415 | 596 | 0.85 | 406 | 529 | 0.79 | 406 | 590 | 0.85 |
| 17 | 415 | 592 | 0.85 | 404 | 569 | 0.79 | 406 | 592 | 0.85 |
| 18 | 415 | 593 | 0.85 | 404 | 521 | 0.79 | 406 | 612 | 0.85 |
| 19 | 415 | 597 | 0.85 | 406 | 573 | 0.79 | 404 | 613 | 0.85 |
| 20 | 415 | 589 | 0.85 | 406 | 570 | 0.79 | 406 | 591 | 0.85 |

**Table S6.** Coefficients of determination (R^2^) and root mean square error (RMSE) between all Chlorophyll parameters and VIs for all samples. Presented are VIs with R^2^ value bigger than 0.5.

|  | **Chl-a (μg cmˉ²)** | | | **Chl-b (μg cmˉ²)** | | | **TChl (μg cmˉ²)** | | |
| --- | --- | --- | --- | --- | --- | --- | --- | --- | --- |
| **Ranking** | **VIs** | **R^2^** | **RMSE**  **(μg cmˉ²)** | **VIs** | **R^2^** | **RMSE**  **(μg cmˉ²)** | **VIs** | **R^2^** | **RMSE**  **(μg cmˉ²)** |
| 1 | NDSI_415, 614_* | 0.86 | 2.23 | NDSI_406, 525_* | 0.79 | 0.65 | NDSI_406, 614_* | 0.86 | 2.74 |
| 2 | ZMI | 0.82 | 2.52 | Carter 1 | 0.64 | 0.86 | GM1/Carter 1* | 0.85 | 2.84 |
| 3 | CIred-edge | 0.82 | 2.52 | TGI | 0.64 | 0.87 | GM1-Carter 1* | 0.83 | 2.96 |
| 4 | Carter 1 | 0.82 | 2.53 | GM1 | 0.64 | 0.87 | Carter 1 | 0.80 | 3.25 |
| 5 | CIgreen | 0.82 | 2.53 | CIgreen | 0.63 | 0.88 | GM1 | 0.80 | 3.27 |
| 6 | GM1 | 0.82 | 2.54 | TCARI | 0.62 | 0.89 | CIgreen | 0.80 | 3.28 |
| 7 | GM2 | 0.81 | 2.56 | Datt1 | 0.59 | 0.92 | ZMI | 0.78 | 3.38 |
| 8 | NDRE | 0.81 | 2.59 | REIP | 0.58 | 0.93 | CIred-edge | 0.78 | 3.38 |
| 9 | REIP | 0.80 | 2.65 | MCARI | 0.58 | 0.93 | GM2 | 0.78 | 3.42 |
| 10 | TGI | 0.79 | 2.74 | NDRE | 0.57 | 0.94 | NDRE | 0.78 | 3.44 |
| 11 | TCARI | 0.78 | 2.76 | ZMI | 0.57 | 0.94 | TGI | 0.78 | 3.45 |
| 12 | MCARI | 0.78 | 2.78 | CIred-edge | 0.57 | 0.94 | REIP | 0.77 | 3.47 |
| 13 | Datt1 | 0.77 | 2.83 | GM2 | 0.57 | 0.95 | TCARI | 0.77 | 3.50 |
| 14 | Ctr2 | 0.76 | 2.89 | Ctr2 | 0.55 | 0.96 | MCARI | 0.76 | 3.59 |
| 15 | SR | 0.75 | 2.97 | NDVI | 0.55 | 0.97 | Datt1 | 0.75 | 3.64 |
| 16 | NDVI | 0.74 | 3.01 | SR | 0.53 | 0.99 | Ctr2 | 0.74 | 3.75 |
| 17 | PRI | 0.65 | 3.48 | - | - | - | SR | 0.72 | 3.86 |
| 18 | Lic2 | 0.64 | 3.52 | - | - | - | NDVI | 0.72 | 3.88 |
| 19 | SIPI | 0.64 | 3.55 | - | - | - | PRI | 0.62 | 4.46 |
| 20 | Lic1 | 0.60 | 3.74 | - | - | - | SIPI | 0.62 | 4.49 |
| 21 | OSAVI | 0.59 | 3.79 | - | - | - | Lic2 | 0.61 | 4.54 |
| 22 | NPCI | 0.51 | 4.15 | - | - | - | Lic1 | 0.58 | 4.73 |
| - | - | - | - | - | - | - | OSAVI | 0.57 | 4.79 |

All R^2^ values are significant to *P* < 0.0001; * marks VIs that were developed in the current study.

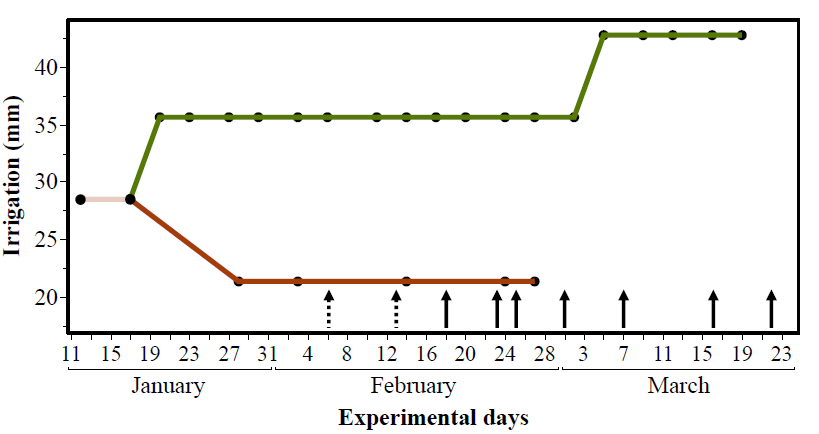

**Figure S1.** Scheme of the experimental design and time point at which chlorophyll samples were analyzed (indicated by arrows). The green line indicates well-watered (WW) regime, and the brown line represents water-limited (WL) regime.

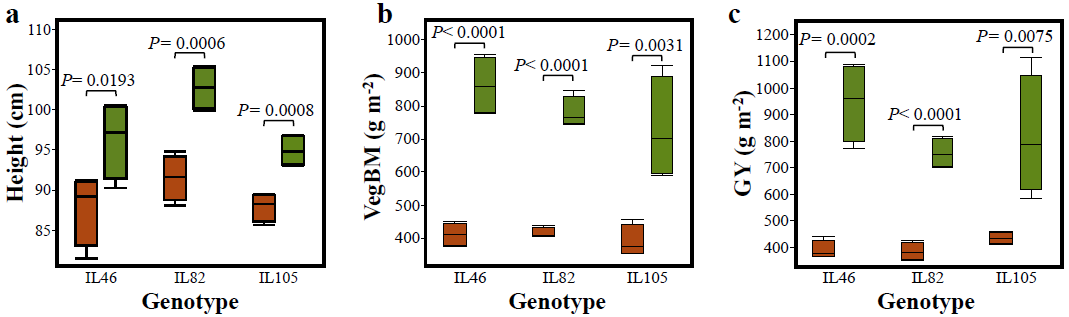

**Figure S2.** Response of the three genotypes to well-watered (WW; green) and water-limited (WL; brown) irrigation regimes. *P*-values indicate a comparison between irrigation regimes using the student t-test. Plant height measured at the end of the season prior to arvest (a). Vegetative biomass (VegBM) obtained at the end of the season by harvest (b). Grain yield (GY) obtained grains obtained at the end of the season by harvest (c).

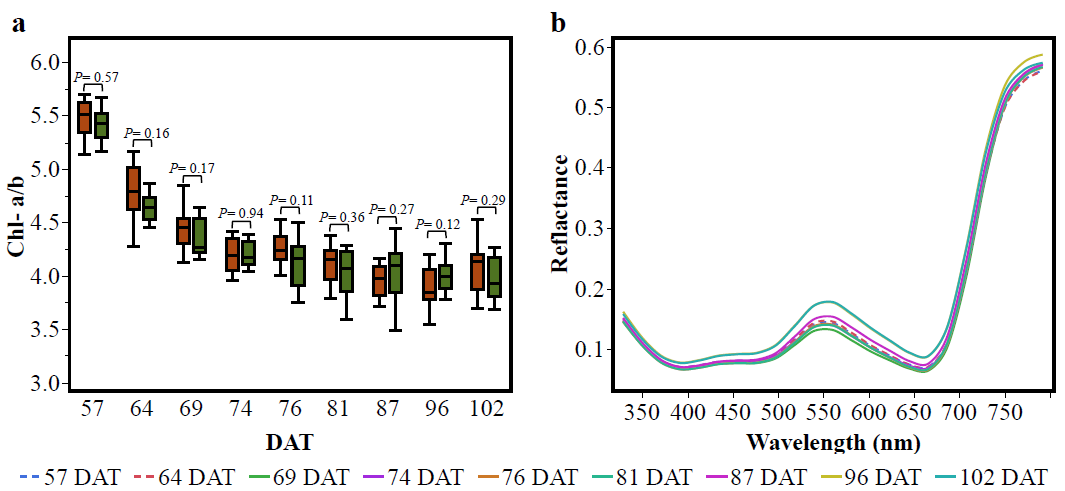

**Figure S3.** )**a**) A Time series of Chl-a/b in response to well-watered (WW; green) and water-limited (WL; brown) irrigation regimes throughout the season. *P*-values indicate a comparison between irrigation regimes using the student t-test. Days after transplant (DAT). (**b**) Spectral reflectance of leaves. Each spectral curve is an average of all leaf spectra acquired that day. The two dates of fully developed leaf spectra (dashed lines) are similar to the flag-leaf spectra throughout the season but in a closer look, they did not follow the chronological intensity trend of the flag-leaf spectra in the visible spectral region and presented the smallest reflectance values around 760 nm but mid-value reflectance around 550 nm.

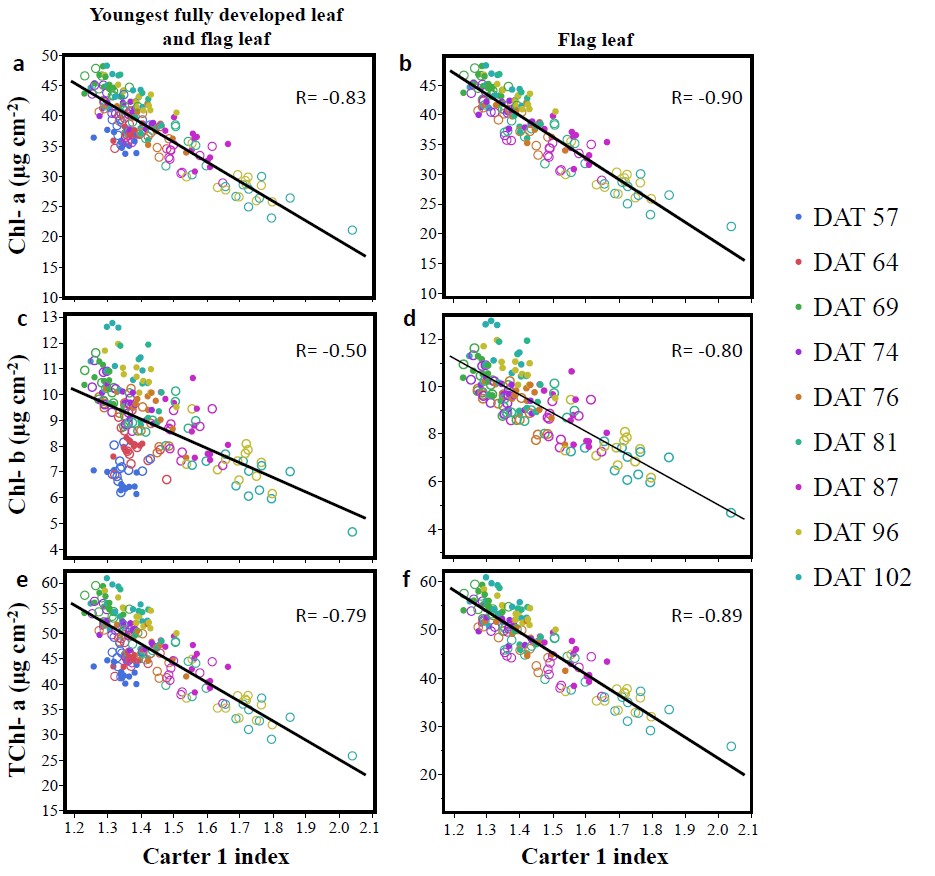

**Figure S4.** Relationship between Carter 1 and Chl-a (**a-b**), Chl-b (**c-d**), and TChl (**e-f**) for data with all dates (a, c, and e), and data from 69 to 102 DAT (b, d, and f). R-values significant at *P*<0.0001. filled stands for well-watered; Hollow stands for water-limited; and DAT stands for dates after transplant.
